# Population genomic evidence of a Southeast Asian origin of *Plasmodium vivax*

**DOI:** 10.1101/2020.04.29.067439

**Authors:** J. Daron, A. Boissière, L. Boundenga, B. Ngoubangoye, S. Houze, C. Arnathau, C. Sidobre, J.-F. Trape, P. Durant, F. Renaud, M.C. Fontaine, F. Prugnolle, V. Rougeron

**Affiliations:** Laboratoire MIVEGEC (Université de Montpellier-CNRS-IRD), CREES, 34394 Montpellier, France; Centre Interdisciplinaire de Recherches Médicales de Franceville, Franceville, Gabon; Service de Parasitologie-mycologie CNR du Paludisme, AP-HP Hôpital Bichat, 46 rue H. Huchard, 75877 Paris Cedex 18, France; Groningen Institute for Evolutionary Life Sciences (GELIFES), University of Groningen, PO Box 11103 CC, Groningen, The Netherlands

## Abstract

*Plasmodium vivax* is the most prevalent and widespread human malaria parasite, with almost three billion people living at risk of infection. With the discovery of its closest genetic relatives in African great apes (*Plasmodium vivax-like*), the origin of *P. vivax* has been proposed to be located in the sub-Saharan African area. However, the limited number of genetic markers and samples investigated questioned the robustness of this result. Here, we examined the population genomic variation of 447 human *P. vivax* strains and 19 ape *P. vivax-like* strains originating from 24 different countries across the world. We identified 2,005,455 high quality single-nucleotide polymorphism loci allowing us to conduct an extensive characterization to date of *P. vivax* worldwide genetic variation. Phylogenetic relationships between human and ape *Plasmodium* revealed that *P. vivax* is a sister clade of *P. vivax-like*, not included within the radiation of *P. vivax-like*. By investigating a variety of aspects of *P. vivax* variation, we identified several striking geographical patterns in summary statistics as function of increasing geographic distance from Southeast Asia, suggesting that *P. vivax* may derived from serial founder effects from a single origin located in Asia.

## Introduction

*Plasmodium vivax* is the most prevalent human malarial parasite responsible for 70 to 80 million clinical cases each year (*1*). It is widespread along the world tropical belt where almost three billion people live at risk of infection. It is the most widely distributed cause of human malaria (*2*). The vast majority of *P. vivax* transmission is located in Central and South East Asia, while populations present in the sub-Saharan African area are protected from transmission due to the absence of the Duffy antigen (*i.e.*, Duffy negativity) at the surface of their red blood cells (*3, 4*). Historically, *P. vivax* remained understudied compared to *P. falciparum* because of its lower mortality rate. However, recent emergence of new therapeutic resistances and the discovery of fatal cases due to *P. vivax* questioned the benign status of *P. vivax* malaria (*5*). In addition, the identification of strains able to invade Duffy-negative red blood cells have been raising concerns over potential spread of the disease into regions that were previously assumed to be protected (*6*). Today, the perception has changed and *P. vivax* is now recognized as a major threat for Public Health (*7*). In order to carry effective strategies for malaria control and elimination, we need to accumulate knowledge on the genetic structure of individual infections, as it provides understanding on the local patterns of malaria transmission and the dynamics of genetic recombination in natural populations of *P. vivax*.

Until recently, early works investigating patterns of genetic variation in worldwide *P. vivax* populations have been limited to a reduced set of samples and genetic markers (based on mitochondrial or autosomal markers) (*8–10*). Consequently, *P. vivax* genetic analyses yielded an incomplete picture of the evolutionary history of this pathogen. Recently, technological improvement has made possible to sequence the whole *P. vivax* genome *via* an enrichment of the parasite DNA from clinical blood samples (*11*). This methodological breakthrough marked the beginning of a new era in the field, with the released in a few years of several projects characterizing the pathogen genetic variation at the whole genome scale in several *P. vivax* populations (*12–15*). Those projects shed light on strong signals of recent evolutionary selection partly due to known drug resistance genes. However, even though they released hundreds of complete genomes, each of them restricted their investigation on a sub-fraction of the worldwide *P. vivax* diversity, either located in Asia-Pacific or South America. Consequently, a comprehensive picture of the worldwide genetic diversity and structure of *P. vivax* populations is still missing and its evolutionary history and how it spread over the world is still poorly understood. Moreover, despite growing evidence suggesting an underlying widespread presence of *P. vivax* across all malaria-endemic regions of Africa (*16*), this continent remains remotely covered with too few complete genomes sequenced from this area. Thus, a key challenge is to provide a worldwide understanding of the genetic variability of *P. vivax* at the genome level to bring insights on the past demographic history and origin of this pathogen.

The origin of current *P. vivax* in humans has stimulated passionate and exciting debates for years. Certain studies placed the origin of the human *P. vivax* in Southeast Asia (“out of Asia” hypothesis) based on its phylogenetic position in a clade of parasites infecting Asian monkeys (*17–19*). This scenario was also supported by genotyping data at 11 microsatellite markers collected across four continents, showing the highest microsatellite diversity in Southeast Asia (*20, 21*). However, the Asian-origin has been challenged by an “out of Africa” scenario, with the recent discovery of a closely related *Plasmodium* species circulating in wild-living African great apes (chimpanzees and gorillas) (*22*). This new lineage, hereafter referred to as *P. vivax-like*, was suggested to have given rise to *P. vivax* in humans following the transfer of parasites from African apes (*23*). This new finding echoes a 70 years-old postulate speculating that the high prevalence of Duffy negativity among human populations in sub-Saharan Africa is a consequence of a long interaction between humans and its parasite, in favor of an African origin of *P. vivax (24*). Yet, despite over a century of research, this monkey’s tale has not yet come to an end. Although now privileged, the “out of Africa” hypothesis remains disputable as the exact series of events liking current human *P. vivax* populations and African great ape *P. vivax-like* still remains unclear (*25*).

Here, to provide novel insight into the worldwide historical demography of *P. vivax* populations, we analyze the genomic variation of 447 human *P. vivax* strains originating from 21 different countries across the world. By including 19 *P. vivax-like* strains sampled on great apes, we explore the evolutionary history of *P. vivax* from its origins to contemporary time and specifically assess whether the origin of the parasite is consistent with an “out-of-Africa” or “out-of-Asia” hypothesis. To our knowledge, this data set provides the most comprehensive characterization to date of the worldwide population genetic structure and diversity of *P. vivax* and *P. vivax-like*, with the identification of two distinct non-recombining *P. vivax-like* clades circulating in sympatry among great apes. Our result clearly demonstrate that *P. vivax* is a sister clade to *P. vivax-like* and is not included within the radiation of *P. vivax-like*, as previously suggested by Liu et al., (*22*). Finally, we discovered multiple lines of evidence from summary statistics of *P. vivax* worldwide genetic variation increasing with geographic distance from Southeast Asia, supporting the hypothesis of a serial founder effect from a single origin located in South-East Asia.

## Results and Discussion

### Genomic data and diversity in *P. vivax and P. vivax-like*

Genome wide data from a total of 1,154 *P. vivax* isolates sampled from all around the world were processed and analyzed (Figure 1A). This dataset included 20 newly sequenced African isolates (Mauritania n=14, Ethiopia n=3 and Sudan n=3), as the worldwide sampling was lacking *P. vivax* isolates coming from Africa (with only three genomes from Madagascar (*13*) and 24 from Ethiopia (*14*)). *P. vivax* samples were obtained from 24 countries across the globe (769 from Asia, 338 from America, and 47 from Africa) (Supplementary Table 1). In order to trace the genetic ancestry of *P. vivax*, we also added a total of 27 genomes of *P. vivax-like* isolated from African great apes. Among them, ten *P. vivax-like* genomes from Gabon were newly sequenced in our laboratory and 17 genomes were obtained from public databases (Gabon n=11, Cameroon n=5 and Ivory Coast n=1) (*23, 25*).

**Figure 1:**
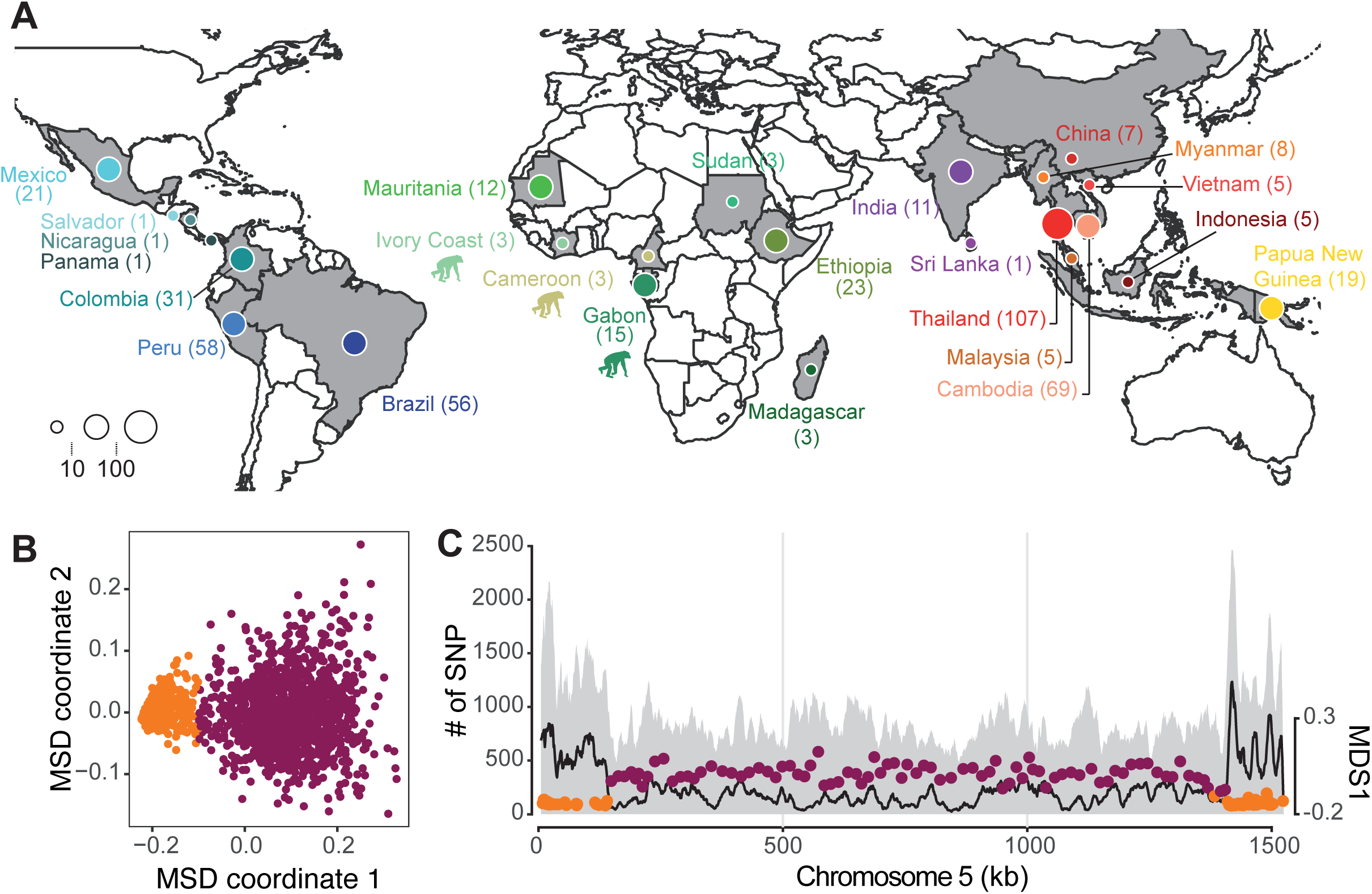
Geographical origin of *Plasmodium* isolates and patterns of genomic variation. A. Country of origin of 447 *P. vivax* and 19 *P. vivax-like*. Note that the isolates are coming from various locations within each country. The chimpanzee pictogram represents African *P. vivax-like* isolates. B. Local variation along the genome in *P. vivax* individual genetic relatedness along the genome visualized with a multidimensional scaling (MDS) based on the local PCA approach (*28*). Each point represents a non-overlapping genomic window of 100 SNPs. Based on the variation of the MDS-1 coordinate, each window has been classified into two groups, the sub-telomeric hyper-variable regions (orange) or the core region (purple). C. Chromosome 5 genome scan of the SNPs count (left y-axis) that has been identified for both *Plasmodium vivax* and *Plasmodium vivax-like.* The black line represents the number of SNPs shared between the two species. The right y-axis represents the MDS-1 coordinates against the middle point of each window. The color represents windows classified within the sub-telomeric hyper-variable regions (orange) or the core region (purple).

Due to the heterogeneity in DNA enrichment methods and sequencing technologies used to access *Plasmodium* genome, our cohort exhibited a broad range of sequencing depth coverage (Supplementary Figure 1). We selected genomes with a minimum average sequencing depth of at least 5x to conduct reliable analyses, thus reducing sampling from 1,154 to 473 for the *P. vivax* isolates and from 27 to 20 for the *P. vivax-like* isolates. Next, we discarded isolates and variants having a proportion of missing data above 50%. Finally, we evaluated for each sample the within-sample parasite infection complexity by calculating the reference allele frequency (RAF) distribution (Supplementary Figure 2). A RAF of 0% or 100% indicates the presence of either the reference or a single alternative allele for all polymorphic sites, which suggests the presence of a single infection (*26*). Overall, nearly all polymorphic variants exhibited either high or low RAF value suggesting the presence of a single infection for all samples. Though, to avoid further possible bias due to multiple infections with several strains, the SNP calling was designed to select within each sample the variant with the highest frequency, resulting in having at each site, one single variant called per sample. After these pre-processing steps, the final data set included a total of 466 genomes (447 of *P. vivax* and 19 of *P. vivax-like*) in which 2,005,455 high-quality SNPs were identified.

### Heterogeneous genetic structure along the genome of *P. vivax*

While population structuration leads to a global impact on the genomic variation in natural populations, local heterogeneity can occur along the genome in regions under the influence of non-random factors, including structural chromosomal features such as large inversions, regions of heterochromatin, or due to selective forces impacting local genetic diversity and recombination (*27*). These factors can lead to local variation in individual strain genetic ancestry and relatedness along the genome confounding global patterns of genetic diversity and population structure. We identified potential local variation in individual relatedness along the genome using a local Principal Component Analysis (PCA) (*28*). As shown in Figure 1B and 1C, contrasted patterns of individual relatedness were found along the genome of *P. vivax*, defining two main genomic partitions: a small partition localized mainly at the sub-telomeric ends of each chromosome (in orange in Figures 1B and 1C), and a larger one at the of each chromosome encompassing 80% of the total SNPs set (hereafter called the “core region”, in purple in Figures 1B and 1C). Interestingly, the sub-telomeric regions identified as having distinct ancestry from the rest of the genome also coincided with hyper-variable genomic regions (in orange in Supplementary Figure 3). Those regions are known to include hyper-variable repetitive regions causing high genotypic errors. Their exact coordinate have only been reported on a former version of the *P. vivax* genome assembly (*12*). The differences in genetic ancestry and strain relatedness provided by the two partitions are evident when looking at the local PCA analyses (Supplementary Figure 4). While the population structure recovered from the core genomic regions displayed interpretable genetic patterns that were consistent with previous studies (*12, 13*), the genetic picture obtained from the SNPs in the sub-telomeric hyper-variable regions was much harder to interpret. We thus excluded these regions from the following analyses, and focused only on the core genomic regions on the central chromosomal region, which represents 21 Mb of the genome and includes 1,610,445 SNPs (∼1 SNP every 13 bp).

### Evolutionary relationships between ape *P. vivax-like* and human *P. vivax*

In order to better understand the evolutionary origin of human *P. vivax*, we examined its phylogenetic relationships with *P. vivax-like* infecting great apes, since chimpanzees and gorillas harbor the closest relatives of human *P. vivax*. The neighbor joining (NJ) tree based on the SNP data set composed of the 19 *P. vivax-like* and 447 *P. vivax* genomes revealed three distinct clades (Figure 2A). The first bifurcation in the tree splits ape-infecting strains from human-infecting strains and represented the strongest axis of genetic variation on the PCA (Figure 2B, 29% of variance explained). This result clearly demonstrates that human *P. vivax* lineage is a sister clade of the ape *P. vivax-like* strains and contradict previous results from Liu et al., suggesting that *P. vivax* forms a monophyletic lineage within the ape parasite radiation (*22*). Liu’s phylogenetic topology, presented as the main argument in favor of an African origin of the parasite, may result from a lack of phylogenetic signal, due to the small number (11 single-genome amplification sequences) of genetic markers investigated. Our result, consistent with previous studies based on a smaller sampling size of *P. vivax-like (23, 25*), re-opens the debate on the origin of *P. vivax* (see below). Interestingly, the second split on the phylogenomic tree (Figure 2A) and on the PCA on the second PC axis (Figure 2B) identified two distinct lineages among *P. vivax-like* strains (referred to PVL.grp1 and PVL.grp2 on Figure 2), composed of five and 14 different strains respectively. Since *P. vivax-like* infects both chimpanzees and gorillas, it would have been reasonable to expect these two lineages to be associated with a specialization on each host. However, both linages were found on both host species.

**Figure 2:**
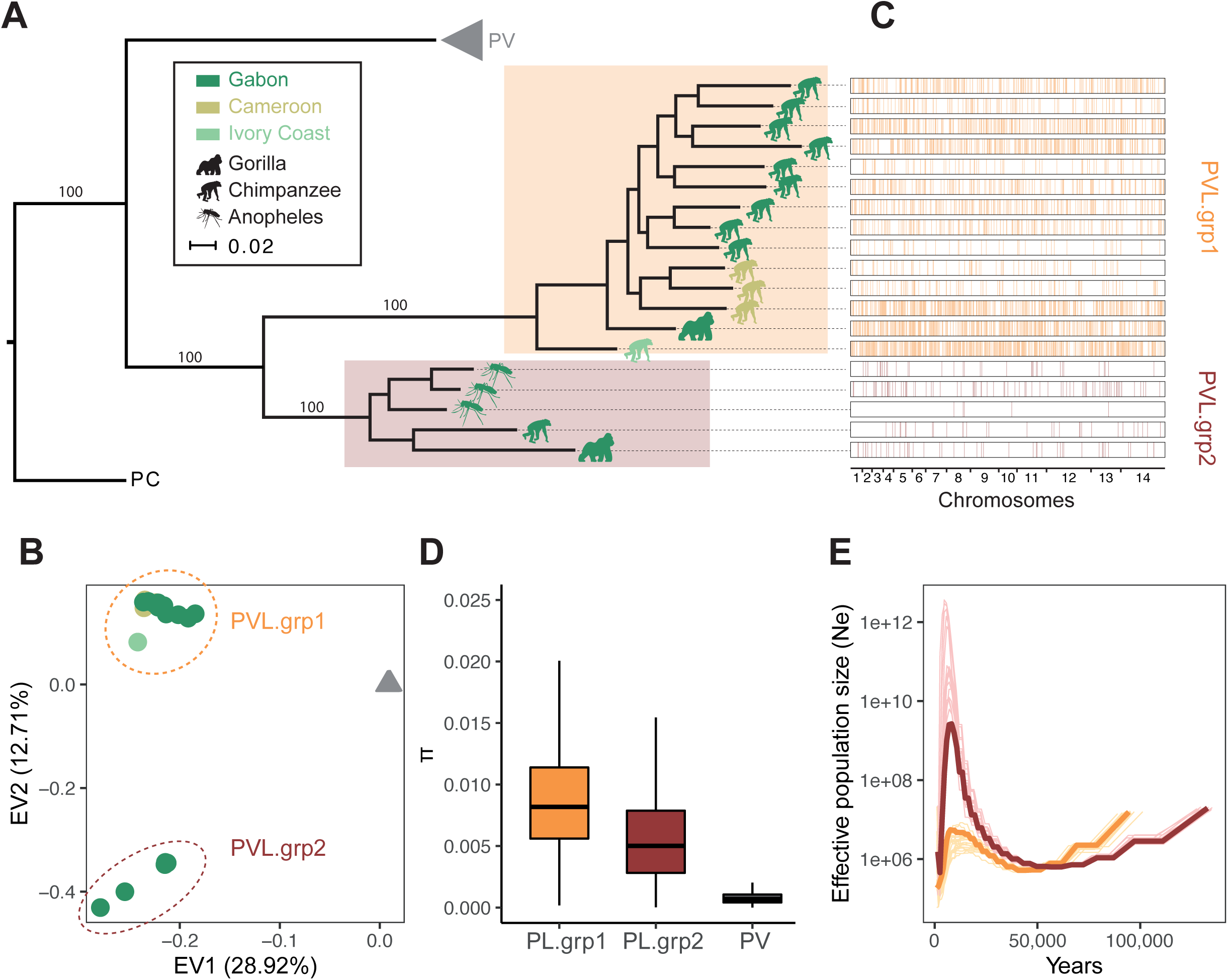
*P. vivax-like* strains are structured into two distinct clades forming a sister monophyletic lineage to the human *P. vivax.* A. Neighbor-joining phylogenetic tree illustrating the relatedness between *P. vivax* (PV, grey triangle) and *P. vivax-like* strains. Based on the phylogeny, two distinct groups of *P. vivax-like* (PVL) were identified: PVL.grp1 (in orange) and PVL.grp2 (in brown). The animal pictograms on each leaf indicate the primate host, gorilla or chimpanzee, colored by country of origin (Gabon, Cameroon, Ivory Coast). The mosquito pictogram represents samples collected on anopheles’ mosquitoes for which the primate host was unknown. The phylogenetic tree was rooted using one outgroup strain of *P. cynomolgi* (strain M). Black dots indicate the bootstrap values, ranging from 0.9 to 1. B. Top two principal components of PCA based on the genome-wide SNPs data present on the core genome of 447 of *P. vivax* and 19 *P. vivax-like* isolates. C. Genome-wide visualization of recent recombination events between the two *P. vivax-like* groups. The horizontal white rectangles represent the genomic position of each *P. vivax-like* individual and the vertical colored lines represent a genomic segment that has recently recombined. The colors indicate donor lineage of each segment. D. Boxplots showing the differences in nucleotide genetic diversity (π) using variants present on the core genome between *P. vivax-like* groups 1 and 2 and *P. vivax.* E. Multiple sequentially Markovian coalescent estimates of the effective population size (Ne) in the *P. vivax-like* groups 1 (in orange) and 2 (in brown) populations. The y-axis shows the log10 of Ne. Light pink and light brown lines represents 50 bootstrap resampling replicates of randomly sampled segregating sites along the core genome.

With the identification of two genetically distinct *P. vivax-like* lineages, we tested the extent of recombination between them, in order to assess whether gene flow still occurs between them. We used the software fastGEAR (*29*) using the 19 *P. vivax-like* individual genomes to identify recently exchanged genomic fragments (*i.e.* those shared between two individuals) and their clade of origin. Through its clustering method, fastGEAR was able to recover the two major *P. vivax-like* clades and identified among them 12,108 recently imported fragments with a mean import length of 1,775 bp (Figure 2C). Interestingly, the donor and recipient individuals of the recently imported fragments belong in all cases to the same lineage, suggesting that no recent inter-lineage recombination occurred between the two distinct *P. vivax-like* clades identified. This result was confirmed further by building a reticulation network for the 19 *P. vivax-like* individuals (Supplementary Figure 5). The lack of reticulation shared between individuals belonging to different lineages is evidence of the absence of recombination between the two distinct *P. vivax-like* clades. Thus, our results clearly demonstrate that although circulating in sympatry inside the same host populations of great apes (*i.e.* within the La Lekedi Park in Gabon), the two lineages of *P. vivax-like* do not recombine, which suggests that they may form distinct species. Intriguingly, a similar subdivision was found in *Plasmodium praefalciparum*, the closest parasite of *P. falciparum*, but the status of these two clades was never investigated as done here for *P. vivax-like (30*).

The genetic diversity (π) of both *P. vivax-like* lineages were found to be about nine higher than the value observed in *P. vivax* (Figure 2D), in agreement with previous reports (*23, 25*). Since it is unlikely that the mutation rates radically differ between *P. vivax-like* and *P. vivax (31*), this difference in genetic diversity may be reflecting a higher historical effective size for *P. vivax-like* than for *P. vivax.* The lower genetic diversity observed in *P. vivax* could result from a bottleneck effect that occurred during the host shift, when the pathogen colonized humans (*23*). A comparable contrasted genetic pattern exists between African apes and humans (*32*), where humans exhibit a far lower genetic diversity than the African apes due to the bottleneck effect that modern human populations underwent with the out-of-Africa expansion from a small number of founders that replaced the archaic forms of humans (e.g. Neanderthals). Is a similar history likely for *P. vivax* in humans? And if yes, does Africa really represent the point of origin of human *P. vivax*, as recently suggested (*22*)? Within *P. vivax-like*, although a higher genetic diversity was detected for PVL.grp1 compared to PVL.grp2, this may reflect sampling differences as PVL.grp1 samples were collected across three African countries, while samples from PVL.grp2 came from one single location, in the Park of La Lékédi in Gabon.

Finally, we inferred historical fluctuations in effective population size (Ne) for the two lineages of *P. vivax-like*. Using a multiple sequentially Markovian coalescent (MSMC) approach, the analysis of genome variation indicated an ancient major expansion in genetic diversity for both clades, follow by a recent dramatic decline (Figure 2E). Unexpectedly, a much high increase in Ne was observed for the PVL.grp2 lineage compared to the PVL.grp1 lineage. Together, our analysis of historical Ne suggest a putative ancient speciation event of the two lineage of *P. vivax-like*, consistent with our previous result demonstrating the reproductively isolation of those two lineages.

### Worldwide genetic structure of human *P. vivax*

To answer these questions, we investigated the genetic structure in our cohort of 447 *P. vivax* isolates collected from 21 countries around the world. Bi-allelic SNPs from the core genome were analyzed using complementary approaches: a PCA that does not rely on any model assumptions (*33*), a model-based individual ancestry analysis implemented in the software ADMIXTURE (*34*), a neighbor joining phylogenetic tree, and by assessing the amount genetic differentiation among pairs of populations using *F*_*ST*_.

All analyses revealed consistent patterns splitting the genetic variation primarily by continents (Figure 3). The first principal component of the genetic variation (EV1) split Southeast Asian from Africa and American populations, while the second axis (EV2) split African populations from the rest of the world (Figure 3A and Supplementary Figure 6). These three major clusters were also identified on the neighbor-joining tree (Figure 3B), and displayed clearly distinct genetic ancestry as estimated when simulating three ancestral clusters (K) by the ADMIXTURE analysis (Supplementary Figure 7).

**Figure 3:**
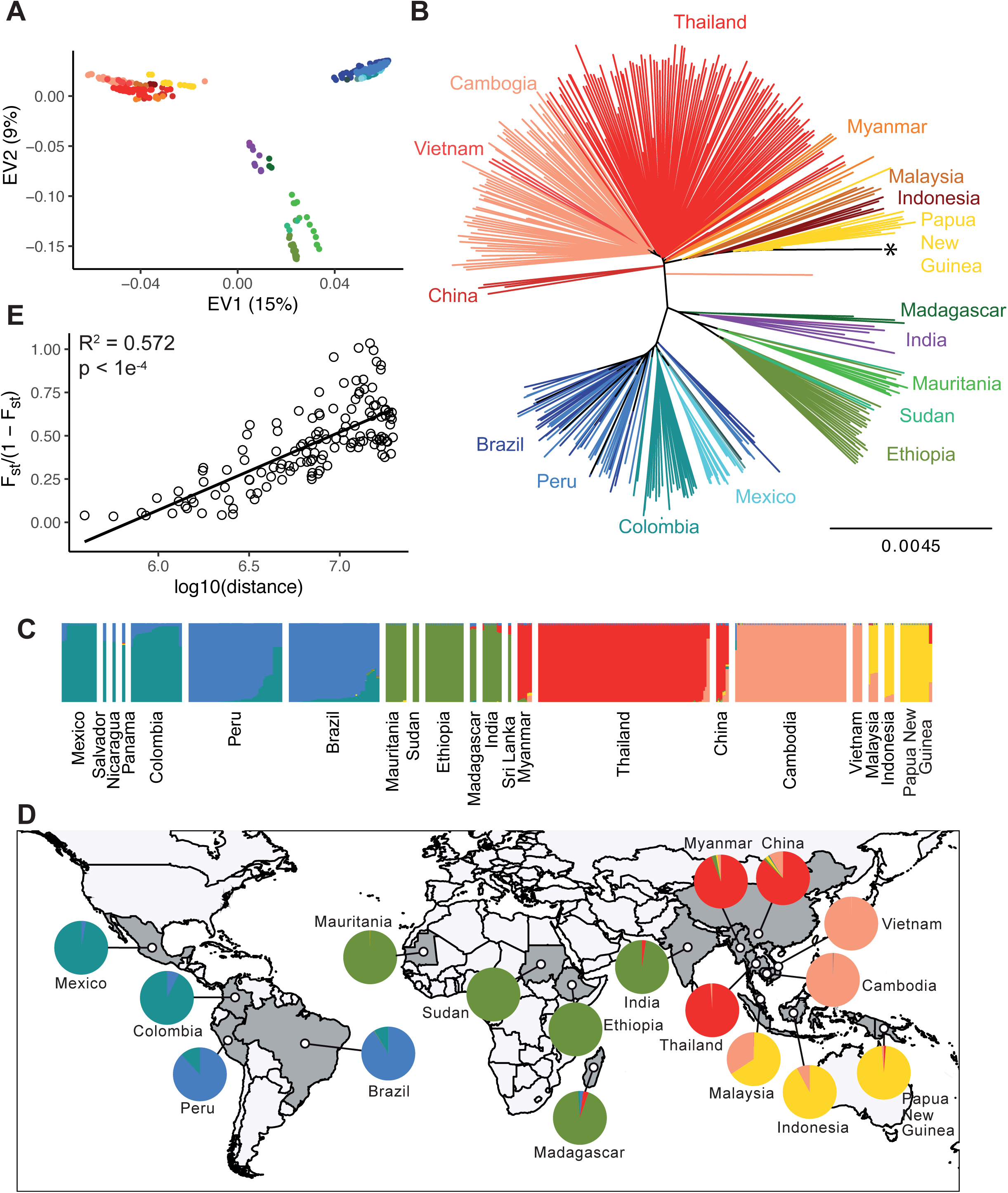
*P. vivax* genetic structure on the core genome is mostly due to genetic isolation of natural populations. A. Principal Component Analysis on the SNPs data from the core genome for the 447 *P. vivax* isolates. B. Neighbour-joining phylogenetic tree constructed using SNPs data present on the core genome. The star indicates the location of the outgroup. C. Isolation by distance among population. Pairwise estimates of *F*_*ST*_ (1-*F*_*ST*_ are plotted against the corresponding geographical distances between countries (as log-10 value). The spearman correlation coefficient and the p-value estimated using a Mantel test with 1000 permutations are shown. D. Individual strain genetic ancestry proportion, which is depicted as a vertical bar, estimated using the program ADMIXTURE (*34*) for each of the K=6 inferred ancestral populations. E. These same ancestral proportions summed over all individuals for each population plotted as pie charts on the world map.

Within each cluster/continent, further subdivision was also evident. Both the PCA (Supplementary Figure 6) and ADMIXTURE (Supplementary Figures 7 and 8) analyses suggested that up to six distinct genetic pools were present. Three distinct ancestral populations were identified in Asia, one in Africa and two in America (Figures 3C and D).

Interestingly, strains from India and Sri Lanka were genetically more closely related to the African populations than to the other neighboring Asiatic groups (Figures 3A, 3B and 3D). This may results from the different trades that took place between the Indian Subcontinent and Africa over the last two millennia, during which movements of populations (and likely diseases) occurred either from India to East Africa or from Africa to India (*35*). The most recent exchange has been the migration of the human Karana population from the north-west India into Madagascar at the end of the 17th century (*36*).

Across the Americas, *P. vivax* strains were structured into two distinct ancestral populations. This is consistent with recent findings suggesting two successive migratory waves responsible for the introduction of *P. vivax* in America (*37*). The first wave has been suggested to occur following a reverse Kon-Tiki route, with a long-range oceanic crossing from the Western Pacific to the Americas. The second wave, has been recently attributed to an introduction by the European colonization of the Americas during the 15th century, and represents the major genetic contributors to the New World *P. vivax* lineages (*38*).

Lastly, individuals collected across the Southeast Asia/Pacific region were structured in three distinct ancestral populations. Among each ancestral population located in Southeast Asia, the genetic differentiation estimated with F_ST_ between countries was weak (Supplementary Figure 9), consistent with relatively unrestricted gene flow between countries. If a distinct ancestral population shared by populations located in Indonesia and Papua New Guinea (PNG) is well explained by their insular isolation, the presence in Southeast Asia continent of two ancestral populations may reflect the effect of the malaria-free corridor previously established through central Thailand, consistent with previous observations in *P. falciparum* (*39*). Population in Malaysia were found admix between two ancestral populations, at a contact zone in which admixture proportions progressively change from one cluster to the other.

### Patterns of genetic diversity in agreement with an Asian origin of human *P. vivax*

The topology of the NJ tree suggested that populations split from each other following a stepping stone colonization model along the world tropical belt. Such stepping stone pattern is expected to lead to an isolation-by-distance (IBD) among populations. As expected under a 2-dimensions stepping stone IBD model (*40*), we found a highly significant relationship between the *F*_ST_/(1-*F*_ST_) and the logarithm of the geographic distance (Mantel test *r*^*2*^= 0.57, p < 10^−4^, Figure 3E), suggesting limited *P. vivax* dispersal across space. Based on the NJ tree, the *P. vivax* clade containing Malaysia, Indonesia and PNG is located nearest to the root of the tree, outward from which are branches that correspond, sequentially to populations from Southeast Asia, Africa and America. Consequently, the branching pattern largely supports an “out-of-Asia” model of *P. vivax* origin rather than the previously suggested “out of Africa” model (*22*).

Under a scenario where the range expansion experienced by *P. vivax* results in a series of founder events, it has been demonstrated that the newly founded populations should experience striking geographical patterns in summary statistics describing *P. vivax* genetic diversity (*41*). First, we investigated the nucleotide diversity (*π*) of our *P. vivax* populations that includes at least five genomes per country, as it is expected to decrease from the native origin to newly invaded areas (*42*). Interestingly, populations from Malaysia and Indonesia displayed the highest genetic diversity, consistent with an Asian origin hypothesis (Figure 4A). We then searched for the hypothetical origin and observed that the decrease in genetic diversity with increasing geographic distance was maximal when Southeast Asia was considered as the putative origin of *P. vivax*, consistent with an Asian origin hypothesis (Supplementary Figure 10). Further evidence for this hypothesis was the very strong correlation we observed considering the putative Asian origin located at the border of Malaysia and Indonesia (spearman *r* = -0.79, p-value < 8×10-4, Figure 4B). By using estimates of the number of cases per year obtained for each country (WHO 2010 report (*43*)), we observed that the correlation between the genetic diversity and distance from Asia remains significant, regardless of the *P. vivax* number of cases (Generalized Linear Models (GLM), p-value < 1.98×10^−5^).

**Figure 4:**
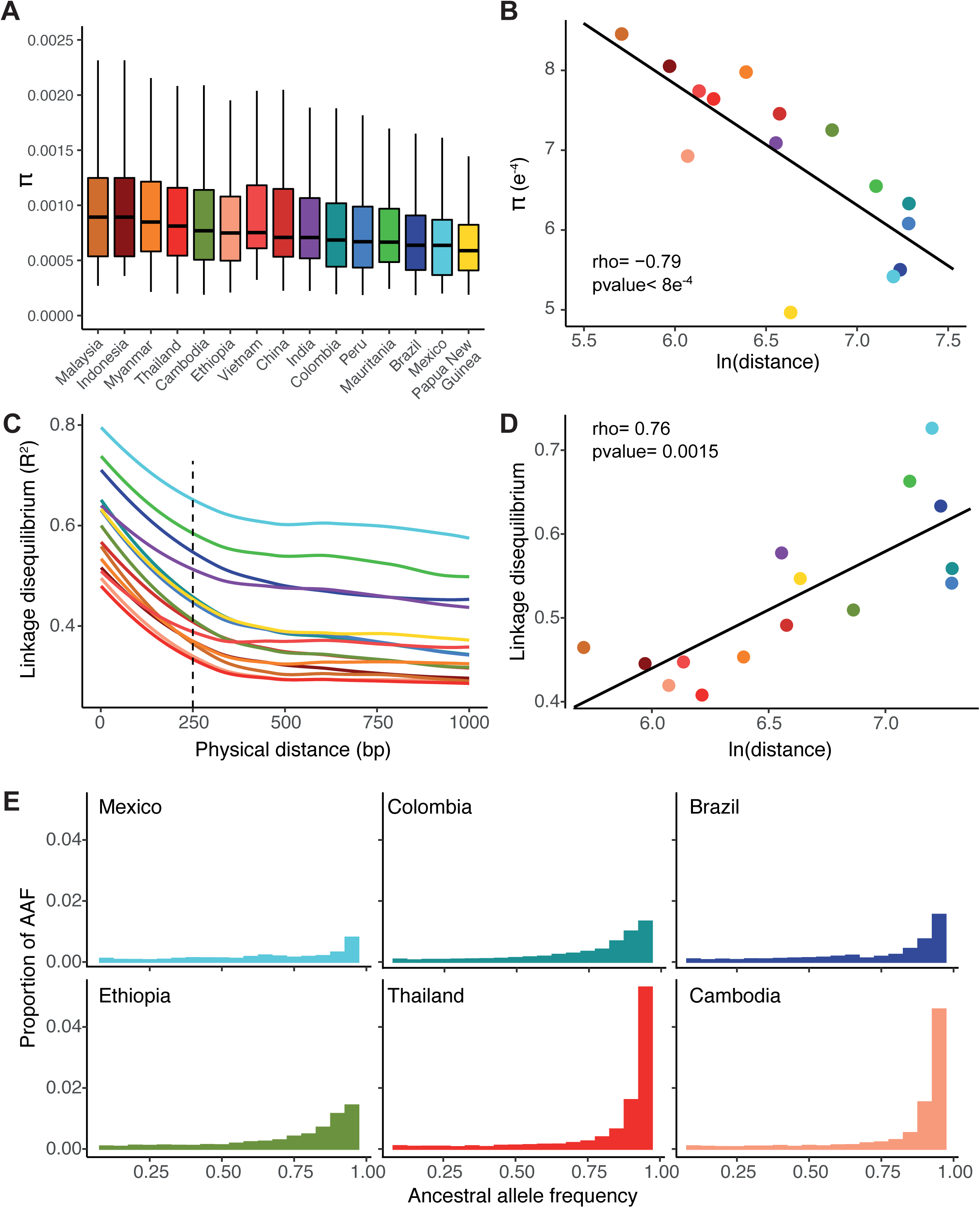
Southeast Asian origin of *P. vivax* is supported by patterns of nucleotide diversity, linkage disequilibrium, and ancestral allele frequency spectrum. A. Boxplot of nucleotide diversity (π) for the different populations of *P. vivax*. П values were calculated at each bi-allelic site among all individuals present in each population. B. Decay of the nucleotide diversity as a function of the log10 distance from the Indonesia-Malaysia border. C. LD-decay of each population of *P. vivax*. LD was measured by R^2^ as a function of physical distance in base pairs (bp). Dotted line represents the physical distance threshold at which the R^2^ values were taken to draw the following regression with the geographic distance. D. LD at 250bp measured by normalized R^2^ as a function of the log10 distance from Malaysia. E. Histograms of the Ancestral Allele Frequency (AAF) in populations with a minimum sample size of 20 individuals are shown. Refer to Figure 1 for the color code that is consistent across the figure.

Next, under a model of sequential founder effects during a range expansion, we would expect linkage disequilibrium (LD) among loci to increase at each step of the expansion, being lowest closed to native populations of origin and highest in populations that are at the colonization front (*44*). We analyzed how LD decay within population varied as a function of the geographic distance from the Asian putative origin identified based on the genetic diversity (Figure 4C). A highly significant and strong positive correlation was observed between LD at 250 bp within each population and the geographic distance to the putative Asiatic origin (rho =0.75, p-value=0.0015, Figure 4D). This result is thus also consistent with a South Asian origin of *P. vivax* that would have spread to the rest of the world.

Third, pattern of ancestral allele frequency (AAF) distributions can also inform on the origin of *P. vivax* and its worldwide colonization routes (*45*). We polarized the AAF spectrum of 94K SNPs in our panel using the *P. vivax-like* and *P. cynomolgi* and considered populations with at least 20 genomes because sampling size can strongly impact the shape of the AAF (Figure 4E). By comparing the shape of the AAF spectrum among populations, we observed that populations located in Southeast Asia displayed more SNPs with high AAFs (>0.9) and fewer with low AAFs, while at the opposite, non-Asian population displayed a progressive flattening of their AAFs distribution. Theoretical work and simulations demonstrated that interplay between multiple demographic forces can have a major role in the change of the AAF spectrum over time (*41*). Large size population would tend to preserve the ancestral state of the variant loci, while small effective population size –that experienced a severe bottleneck-would have more pronounced genetic drift, resulting in a more rapid increase in derived allele frequencies. Our results based on the AAFs spectrum show that population from Africa displayed a lower amount of ancestral allele than population from Asia, consistent with a South Asian origin of *P. vivax*.

Finally, we analyzed the evolution of the Ne through time and their time to the most recent common ancestor (TMRCA) for populations belonging to the different geographical regions. In a serial founder-colonization model, the TMRCA is expected to decrease with distance from the origin, as observed for humans (*46*). To test this hypothesis, we applied the multiple sequentially Markovian coalescent (MSMC) approach to perform demographic inference using five individuals from 14 *P. vivax* populations. As previously describe in Otto et al. (*31*) we assumed a mutation rate (*µ*) per generation (*g*) of *µ*.*g* = 1.158 × 10^−9^ and a generation time of g = 0.18. As expected, our results show that the TMRCA values in the populations from Southeast Asia were in most of the case older than the TMRCA values of the populations from Africa and America (Figure 5). In addition, the MSMC curves revealed that all populations displayed two distinct dynamics in their effective population sizes. Populations from Southeast Asia experienced an increase in their effective population size in a more distant past, followed by a steady decline. In contrast, populations from Africa and America exhibited only a decrease in their effective population size, with a severe bottleneck in a recent past.

**Figure 5:**
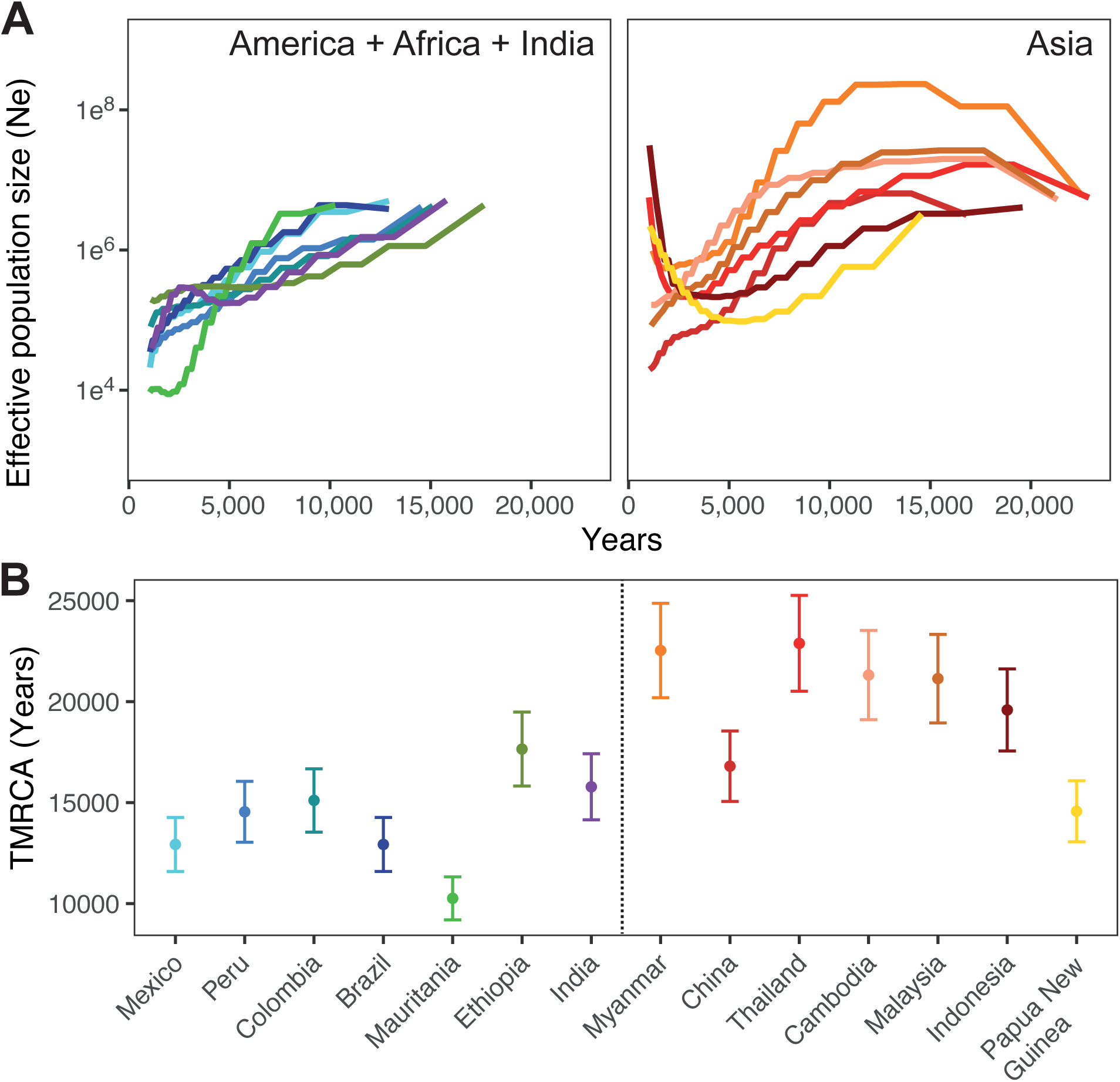
Coalescent-based inference of demographic history in each *P. vivax* population. (A) Effective population size variation back to the time to the most recent common ancestor (TMRCA) inferred using MSMC and (B) inferred TMRCA in each population. Analyses were conducted considering 5 individuals in each population, assuming a mutation rate per generate (*µ.g* = 1.158 × 10^−9^) and a generation time (g) of 0.18.

Together, these results of *P. vivax* populations displaying a decrease in genetic diversity and increase in LD with increasing geographic distance from Southeast Asia are consistent with a model of serial founder events beginning from an origin located in Southeast Asian, more precisely near Malaysia and Indonesia. The higher proportion of ancestral alleles and older TMRCA found in the Southeast Asian populations are additional evidence supporting this scenario. Therefore, these results are more consistent with a scenario in which the ancestral populations that gave rise to the current human *P. vivax* originated from Asia.

An Asian origin from a non-human primate raises the question about the fixation of the Duffy negativity in sub-Saharan Africa as a result of an ancient presence of *P. vivax* on that continent. If it is well admitted that the fixation of the FY*0 allele in sub-Saharan Africa constitutes one of the fastest known selective sweep for any human gene (*47*). Given the relatively mild clinical symptoms of modern *P. vivax*, it is unlikely that the evolutionary forces that led to the allele fixation have been triggered by *P. vivax* alone (*48*). Especially in regard of growing evidence showing the presence of endemic *P. vivax* circulating in sub-Saharan Africa (including Duffy negative hosts), suggesting that the FY*0 allele may confer only partial protection against *P. vivax (16*).

In conclusion, this study provides to our knowledge one of the most detailed view of the worldwide distribution of the population genetic diversity and demographic history thus far available. The present study focused intentionally on the global patterns of genetic diversity in order to assess the origin of *P. vivax*, addressing especially the expectation under the different hypotheses. We showed here that *P. vivax* is a sister group to *P. vivax-like*, and not a sub-lineage. The genetic diversity in *P. vivax-like* is clearly richer than *P. vivax*, consistent with a strong bottleneck in the lineage that gave rise to *P. vivax*. Our results based on whole genome sequencing support of an out-of-Asia origin, rather than Africa, for the world populations of *P. vivax*, with a clear signal of stepping stone colonization events accompanied by serial founder effects. A specific effort should now be done to describe in more details the demographic and selective history of the parasite in some particular regions (e.g. South America, Africa, Europe) and estimate when and how these different regions were colonized. The question of the host of origin still remain open.

## Material and Methods

### African *P. vivax* samples collection and ethical statements

Since only 27 human *P. vivax* genomes from the African continent were publicly available, we augmented this dataset by sequencing 20 additional *P. vivax* genomes from Mauritania (N=14), Ethiopia (N=3) and Sudan (N=3). *P. vivax* infections were diagnosed using microscopy, PCR on the *Cytochrome b* gene and / or Rapid Detection Test (RDT). Samples were collected from *P. vivax*-infected patients after obtaining informed consent and following ethical approval at the local institutional review board of each country. The informed consent procedure for the study consisted of a presentation of the aims of the study to the community followed by invitation of adult individuals for enrolment. At the time of sample collection, the purpose and design of the study was explained to each individual and a study information sheet was provided before oral informed consent was collected. Oral consent was provided since the study did not present any harm to subjects and did not involve procedures for which written consent is required. The verbal consent process was consistent with the ethical expectations for each country at the time of enrolment, approved by each country and the ethics committees approved these procedures. Privacy and confidentiality of the data collected were ensured by the anonymization of all samples before the beginning of the study. For samples from Mauritania, the study received the approval from the pediatric services of the National Hospital, the Cheikh Zayed Hospital and the Direction régionale à l’Action Sanitaire de Nouakchott (DRAS)/Ministry of Health in Mauritania. No ethics approval number obtained at this time. For samples from Sudan, no specific consent was required, the human clinical, epidemiological and biological data were collected in the CNRP database and analyzed in accordance with the common public health mission of all French National Reference Centers (https://www.legifrance.gouv.fr/affichTexte.do?cidTexte=JORFTEXT000000810056&dateTexte=&categorieLien=id). The study of the biological samples obtained in the medical care context was considered as non-interventional research (article L1221-1.1 of the French public health code) only requiring the non-opposition of the patient during sampling (article L1211-2 of the French public health code). All data collected were anonymized before analyses. For samples from Ethiopia, this study was approved by the Ethical Clearance Committee of Haramaya University-College of Health and Medical Sciences, and from the Harari and Oromia Regional State Health Bureau in Ethiopia.

### *P. vivax-like* samples collection and ethical statements

*P. vivax-like* samples were obtained through a continuous survey of great ape *Plasmodium* infections carried out in the Park of La Lékédi, in Gabon, in collaboration with the Centre International de Recherches Médicales de Franceville (CIRMF). The park of La Lékédi is a sanctuary for ape orphans in Gabon. During sanitary controls, blood samples were collected and treated just after collection on the field using leukocyte depletion by CF11 cellulose column filtration (*49*), before being stored at -20°C at the CIRMF. The animals’ well-being was guaranteed by the veterinarians of the “Parc of La Lékédi” and the CIRMF, who were responsible for the preceding sanitary procedures (including blood collection). All animal work was conducted according to relevant national and international guidelines. These investigations were approved by the Government of the Republic of Gabon and by the Animal Life Administration of Libreville, Gabon (no. CITES 00956). It should be noted that our study did not involve randomization or blinding.

*P. vivax-like* samples were also obtained from sylvatic anopheles mosquitoes trapped with CDC light traps in the forest of the same park, during a longitudinal study (*50*). Anopheles mosquitoes were morphologically identified by reference to standard keys (*51*), stored in liquid nitrogen, returned at the CIRMF and kept at –80 °C until analysis.

Genomic DNA was extracted from each sample using DNeasy Blood and Tissue kit (Qiagen, France) according to manufacturer’s recommendations. *P. vivax-like* samples were identified by amplifying and sequencing either *Plasmodium* Cytochrome b (*Cytb*) or Cytochrome oxidase 1 (*Cox1*) genes as described elsewhere (*18, 52*). This resulted in a total of 10 *P. vivax-like* samples to sequence, including 3 from gorillas, 4 from chimpanzees and 3 Anopheles mosquitoes.

### African *P. vivax* and *P. vivax-like* genomes sequencing

To overcome host DNA contamination, selective whole genome amplification (sWGA) was used to enrich submicroscopic DNA levels as already described elsewhere (*53*). This technic preferentially amplifies *P. vivax* and *P. vivax-like* genomes from a set of target DNAs. For each sample, the DNA amplification was carried out by the strand-displacing phi29 DNA polymerase and two sets of *P. vivax*-specific primers that target short (6 to 12 nucleotides) motifs commonly found in the *P. vivax* genome (set1920 and PvSet1) (*52, 53*). For each set of primers separately, 30 ng of input DNA was added to a 50 µl reaction mixture containing 3.5 µM of each sWGA primers, 30 units of phu29 DNA polymerase enzyme (New England Biolabs), 1X phi29 buffer (New England Biolabs), 4 mM of dNTPs (Invitrogen), 1% of Bovine Serum Albumine and sterile water. Amplifications were carried out in a thermal cycler: a ramp down from 35°C to 30°C (10 min per degree), 16h at 30°C, 10 min at 65°C and hold at 4°C. For each sample, the two amplifications obtained (one per set) were purified with AMPure XP beads (Beckman-Coulter) at a 1:1 ratio according to the manufacturer’s recommendations and pooled at equimolar concentrations. Finally, each sWGA pooled library products were prepared using the Nextera XT DNA kit (Illumina) according to the manufacturer’s protocol. Samples were then pooled and clustered on the HiSeq-2500 sequencer in Rapid Run mode with 2×250-bp paired end reads.

### Genomic data from public and/or published data archives

To provide a worldwide comparative study of the *P. vivax* genetic diversity, population structure and evolution, we performed a large literature search to identify previously published genomic data set. We identified and collected fastq files from 1,134 *P. vivax* isolates obtained from the following 12 different bioprojects: PRJEB10888 (*14*), PRJEB2140 (*12*), PRJNA175266 (*54*), PRJNA240356 (*13*), PRJNA240452, PRJNA240531, PRJNA271480 (*13*), PRJNA284437 (*55*), PRJNA350554 (*56*), PRJNA432819 (*57*), PRJNA432819 (*58*), PRJNA65119 (*59*). In addition, 17 *P. vivax-like* sequenced genomes from Cameroon, Ivory Coast and Gabon were collected from two different bioprojects (PRJNA474492 and PRJEB2579) (*23, 25*). Published genome were downloaded from the National Center for Biotechnology Information (NCBI) Sequence Read Archive (SRA) (*60*) and converted into fastq files using the NCBI SRA-Toolkit (*fastq-dump* –split-3). Details about the genomic samples are provided in Supplementary Table 1.

### *P. vivax* and *P. vivax-like* read mapping and variant calling

Both newly generated and previously published sequencing reads were trimmed for adapters and pre-processed to remove low quality reads (--quality-cutoff=30) using the program *cutadapt* (*61*). Reads shorter than 50bp and containing ‘N’ were discarded (--minimum-length=50 --max-n=0). Sequenced reads were aligned to the PVP01 (*62*) *P. vivax* reference genome using *bwa-mem* (*63*). Here, a first round of filtration was applied to our isolates by excluding isolates having an average genome coverage depth lower than 5x. We used the *Genome Analysis Toolkit* (GATK, version 3.8.0) (*64*) to identify SNPs in each isolate following GATK best practices. Duplicate reads were marked using the *MarkDuplicates* tool from the Picard tools (*65*) with default options. Local realignment around indels was performed using the *IndelRealigner* tool from *GATK*. Variant calling was carried out using *HaplotypeCaller* module in *GATK*, on reads mapped with a “reads minimum mapping quality” of 30 (-mmq 30) and a minimum base quality > 20 (--min_base_quality_score 20). We filtered out low quality SNP according to GATK best-practice recommendations and kept SNP call at site covered by at least 5 reads. Finally, individual isolate VCF files were merge using the *GATK* module *GenotypeGVCFs*. Because of our tolerant cutoff of genome depth of coverage set up at 5x, we apply a second round of filtration to our data by excluding variants and isolates with a missing call rate >50%. To evaluate the possibility of co-infections we follow the method developed by Chan et al. (*26*) and examined the reference allele frequency distributions (RAF). Single infection would display a strict RAF pattern as all variants will carry either the reference or a single alternate allele (*i.e* RAF of 100% or 0%). For each of the samples, we calculated the proportion of reads carrying the reference allele to display the RAF distribution.

### Heterogenous patterns of relatedness along *Plasmodium* genomes

In order to identify heterogenous patterns of relatedness along Plasmodium genomes, we conducted a local-PCA analysis, as described in Li et al., 2018 (*28*). Briefly, the *P. vivax* genome was divided into 1,439 contiguous and non-overlapping windows of 100 SNPs. On each window, we applied a principal component analysis (PCA) and stored the individual isolate scores for the first two principal components. To measure the similarity among windows, a Euclidian distance matrix is computed among windows based on the PC scores. Finally, a multidimensional scaling (MDS) was used to visualize the relationships among windows. We used a set of three coordinates to visualize the patterns of relatedness shared among windows. Because the local PCA analysis was sensitive to missing data, we selected the top 304 *P. vivax* genomes having a SNPs missing discovery rate lower than 25% to run this analysis.

### Analysis of genetic recombination among *P. vivax-like* genomes

Recombination among *P. vivax-like* strains was inferred using fastGEAR software (*29*). From the SNPs present in the core genome we created a multi-fasta file used as input for fastGEAR that we launched with the iteration number set to 15 (default). Recent recombination events were detected with the Bayesian factor (BF) > 1 (default) and refers to inter-lineage recombination for which the donor-recipient relation can be inferred. Phylogenetic network tree was generated using SplitTree 4 software (*66*) using the polymorphic sites present on the core genome.

### Population structure analysis

Genetic relationships between populations and species were assessed and visualized using a distance based NJ trees, PCA and by *ADMIXTURE* analyses. PCA and *ADMIXTURE* analyses were performed after selecting only bi-allelic SNPs present in the core *P. vivax* genome, excluding singleton from the dataset. The variants were linkage-disequilibrium-pruned to obtain a set of unlinked variants using the option --indep-pairwise 50 5 1.5 in PLINK (*67*). PCA analysis was performed using PLINK and display in R. Assessment of population structure and estimation of isolate individual ancestry to various number (K) of ancestral populations was performed using *ADMIXTURE (34*). Each ADMIXTURE analysis was repeated 100 times with different random seeds, with a K value ranging from 2 to 20. The most likely number of ancestral populations (*K*) was determined using the cross-validation error. We then used *CLUMPAK (68*) to analyze ADMIXTURE outputs and compute ancestry proportions. NJ tree was estimated using the *ape* R package (*69*) using the full SNP dataset present on the core *P. vivax* genome (without performing any SNP LD-pruning) and plotted with figtree (*70*). The reliability of branching order was assessed by performing 1,000 bootstrap replicates. In order to use *P. cynomolgi* (strain M) as outgroup we used the tools genome liftOver (*71*) to translate our SNP coordinates from the *P. vivax* assembly to the *P. cynomolgi* assembly. Genetic differentiation among populations was estimated using the Weir and Cockerham’s estimator of *F*_ST_ using *VCFtools* (*72*). This metric accounts for differences in the sample size in each population.

### Demographic history analysis

To investigate the demographic history of *P. vivax* and test distinct scenarios, we compute several statistics that describing different features of genetic diversity within populations. Each of these analyses was computed for each of the populations defined by country of origin using the SNPs present on the core genome of the reference. To minimize the impact of missing data on our analyses only individuals with less than 5% of missing data were considered for the following analysis. The genome-wide nucleotide diversity (π) was calculated by averaging the number of nucleotide differences between pairs of DNA sequences at each SNP position using VCFtools. LD-decay was estimated by randomly sampling five individuals from each population using the tools PopLDdecay (*73*), that calculate the genotype correlation coefficient R2 for pairs of SNPs at a maximum range of distances of 10kb. Finally, we polarized the bi-allelic SNPs of each *P. vivax* population using shared alleles identified between *P. vivax-like* individuals and *P. cynomolgi*. The final unfolded SFS was computed for a total of 93,652 SNPs. Regression of nucleotide diversity against geographic distance were calculate as described in Fontaine et al., (*74*). Briefly, we divided the world map into a 400 × 500 pixel two-dimensional lattice, and considered each pixel as a potential source for the geographic expansion of *P. vivax*. The geographic distances between each population and the focal pixel were determined using the R package *geosphere*, to then calculate the spearman coefficient of correlation between the nucleotide diversity and the geographic distance. The pixel with the lowest negative correlation coefficient is thus indicative of the origin.

### Estimates of the effective population size

To infer historical changes in effective population size (*Ne*) and estimate the TMRCA in the *P. vivax* and *P. vivax-like* populations, we ran the MSMC program (*75*) on the core genome of all chromosomes. Homozygous SNPs present on each *Plasmodium* chromosome were considered as single phased haplotype. We run MSMC for 20 iterations with a fixed recombination rate. The MSMC method was applied by selecting randomly five individuals from each population with a missing discovery rate lower than 5%. The error around our estimates was estimated by bootstrapping 50 replicates by randomly resampling from the segregating sites used.

## Supplementary Materials

**Supplementary Table 1**: Genome statistics and metadata. Country of origin, GeneBank accession number and several *in silico* results for the 1,154 human *P. vivax* and 27 *P. vivax-like* strains analyzed in this study.

**Supplementary Figure 1**: Distribution of the mean sequencing depth of *P. vivax* and *P. vivax-like* whole-genome sequence. Genomes with a sequencing depth ≥5x (dash line) were selected for downstream analysis.

**Supplementary Figure 2**: Rare allele frequencies distribution. For each population of *P. vivax* (PV) and *P. vivax-like* (PVL), the distribution of polymophic sites (y-axis) is represented as a function of the proportion of reads carrying the reference allele (x-axis). Almost all polymophic sites, are supported by reads carrying either the reference or a single alternate allele, suggesting the presence of single infection.

**Supplementary Figure 3**: Genome scan of the SNP density (left y-axis) along the 14 chromosomes for *P. vivax* and *P. vivax-like* combined. The black line represents the number of SNPs shared between the two species. The right y-axis represents the scores along the first axis of the multidimensional scaling analysis (MDS1) for each genomic window along the genome.

**Supplementary Figure 4**: Scatter plots showing the results of the two PCAs conducted using the SNPs located either in the core genome or in the hypervariable sub-telomeric regions.

**Supplementary Figure 5**: Reticulate network based on SplitTree4 (*66*), drawn from 19 *P. vivax-like* genomes. The reticulate networks split the *P. vivax-like* genomes in two distinct clades. On the tree, reticulation indicates likely occurrence of recombination. No reticulation was observed among the two clades suggesting the absence of recombination between the two clades.

**Supplementary Figure 6**: PCA analysis of the 447 of *P. vivax* using SNPs present in the core genome. Scatter plot shows individual strain relationships along the principle components 3 to 6. Dot colors indicate populations. The bar chart shows the percentage of variance explained by each principal component axis.

**Supplementary Figure 7**: Individual ancestry proportion to each ancestral population tested (from K=2 to K=10) estimated using the ADMIXTURE program for the 447 P. vivax genomes. Individual strains are sorted by country with increasing longitude.

**Supplementary Figure 8**: Cross validation error rate estimated using the ADMIXTURE program for each ancestral populations tested (K between 2 and 20). We chose K=5 to analyze the SNPs data, as the value that minimizes the error.

**Supplementary Figure 9**: Average allele frequency differentiation estimated using *F*_*ST*_ values between pairs of populations.

**Supplementary Figure 10**: Genetic diversity of *P. vivax* regressed on geographic distance across the world. The value at each pixel of the map corresponds to the Spearman correlation coefficient (*r*) between the expected genetic diversity in each population and the geographic distance between this population and the pixel. The black dots represent the sampling sites used in the regression (where n ≥ 5 individuals).

**Supplementary Figure 11**: Multiple sequentially Markovian coalescent (MSMC) estimates of the effective population size (*Ne*) in 14 *P. vivax* populations. The y-axis shows the log10 of Ne. Gray lines represent the MSMC results obtained from 50 bootstrap resampling of the segregating sites.

## Acknowledgements

**Acknowledgements:** The authors acknowledge the IRD itrop HPC (South Green Platform) at IRD Montpellier for providing HPC resources that have contributed to the research results reported within this paper. URL: https://bioinfo.ird.fr/-http://www.southgreen.fr.

## Author contributions

Conception: V.R. and F.P.; funding acquisition: V.R. and J.D.; biological data acquisition and management: V.R., A.B., B.N., B.L., S.H., C.A., C.S., J-F.T., P.D.; sequence data acquisition: A.B. and V.R.; method development and data analysis: J.D. and M.C.F.; interpretation of the results: J.D., M.C.F., F.P., and V.R.; drafting of the manuscript: J.D. and V.R.; reviewing and editing of the manuscript: J.D., F.R., M.C.F., F.P., and V.R.

## Data and materials availability

The Illumina sequence reads generated on the new *P. vivax* samples from Mauritania, Sudan and Ethiopia have been deposited in the European Nucleotide Archive (ENA) under the accession codes listed in Supplementary table 1. The authors declare that all other data supporting the findings of this study are available within the article and its Supplementary Information files, or are available from the authors upon request.

