## Supplemental Figures and Table for "Population genomic evidence of a Southeast Asian origin of *Plasmodium vivax*": supplementaryFigures.pdf

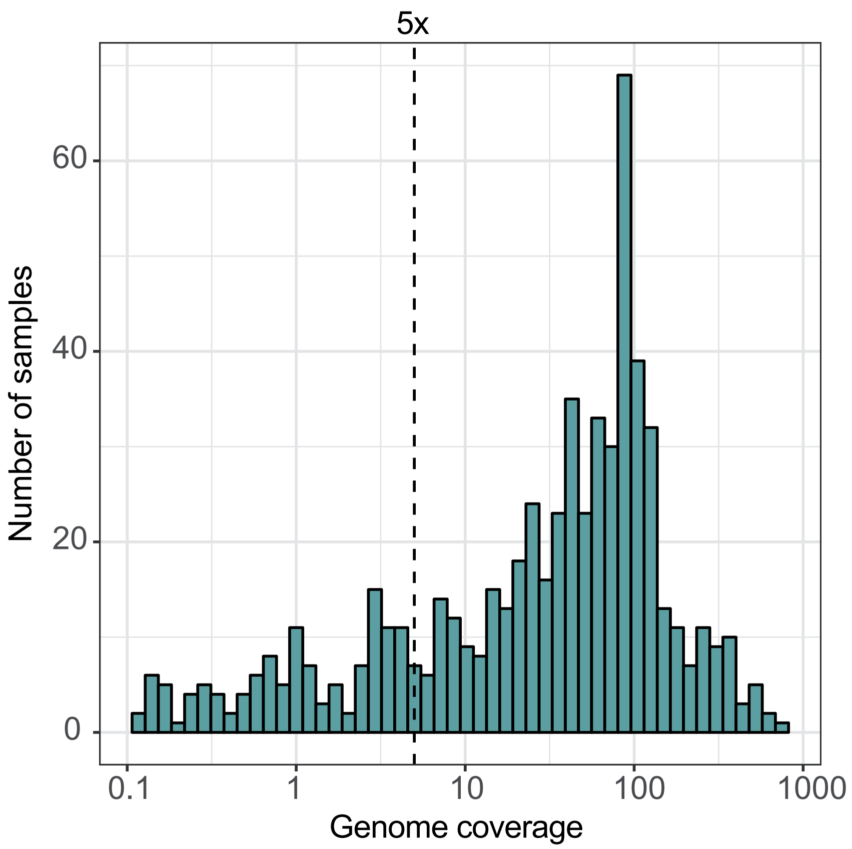


**Supplementary Figure 1:** Distribution of the mean sequencing depth of *P. vivax* and *P. vivax-like* whole-genome sequence. Genomes with a sequencing depth ≥5x (dash line) were selected for downstream analysis.

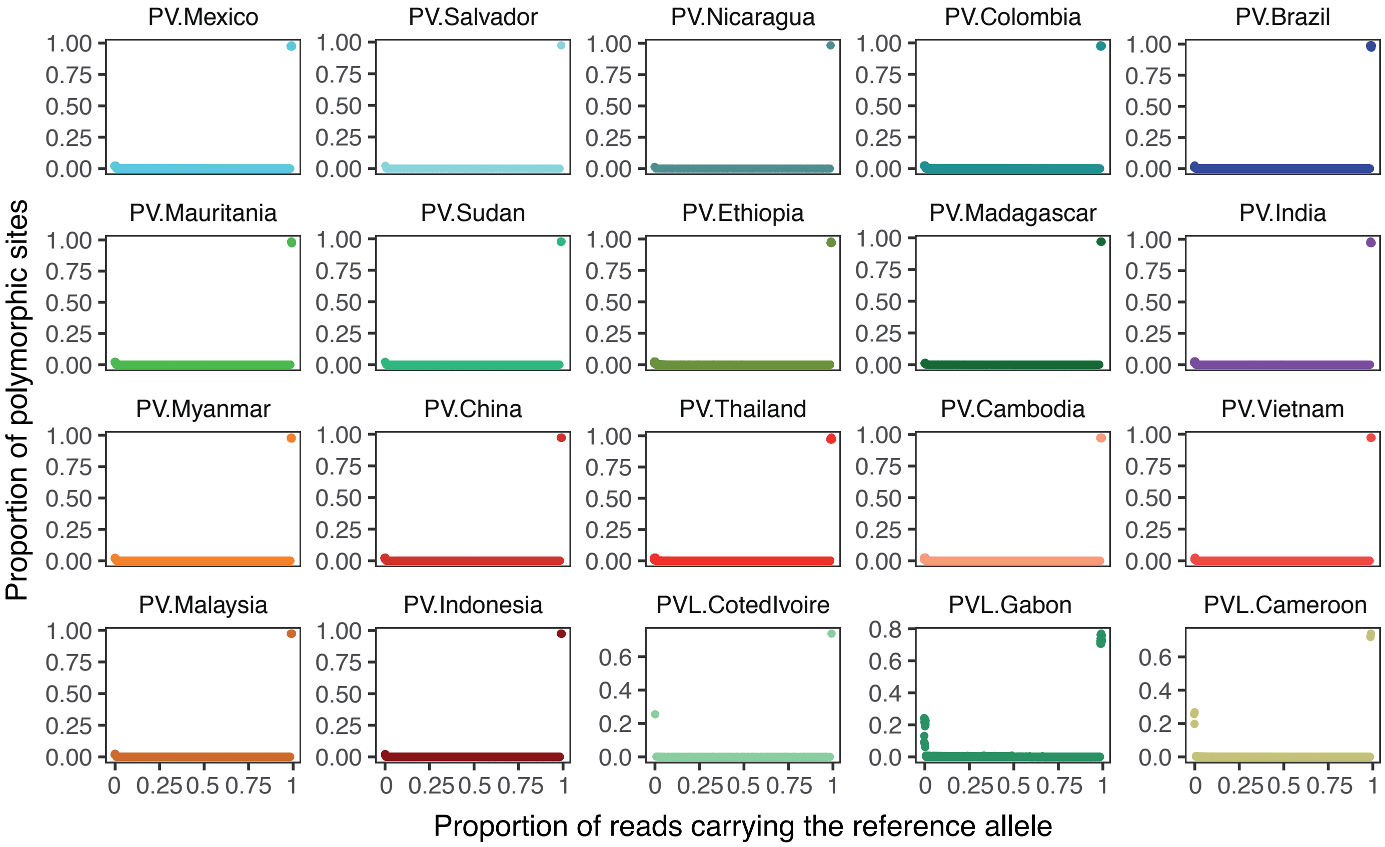


**Supplementary Figure 2:** Rare allele frequencies distribution. For each population of *P. vivax* (PV) and *P. vivax-like* (PVL), the distribution of polymophic sites (y-axis) is represented as a function of the proportion of reads carrying the reference allele (x-axis). Almost all polymophic sites, are supported by reads carrying either the reference or a single alternate allele, suggesting the presence of single infection.

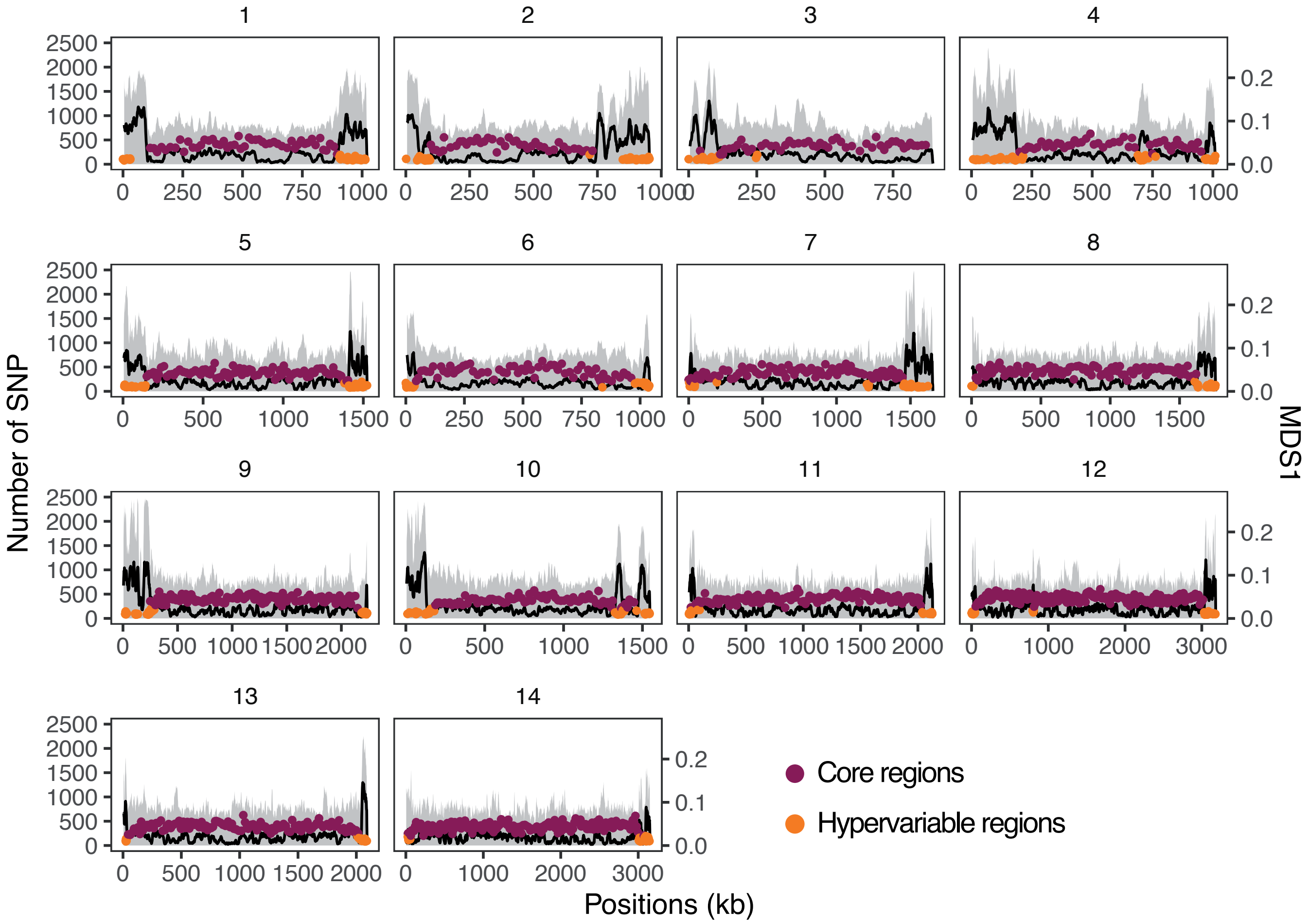


**Supplementary Figure 3:** Genome scan of the SNP density (left y-axis) along the 14 chromosomes for *P. vivax* and *P. vivax-like* combined. The black line represents the number of SNPs shared between the two species. The right y-axis represents the scores along the first axis of the multidimensional scaling analysis (MDS1) for each genomic window along the genome.

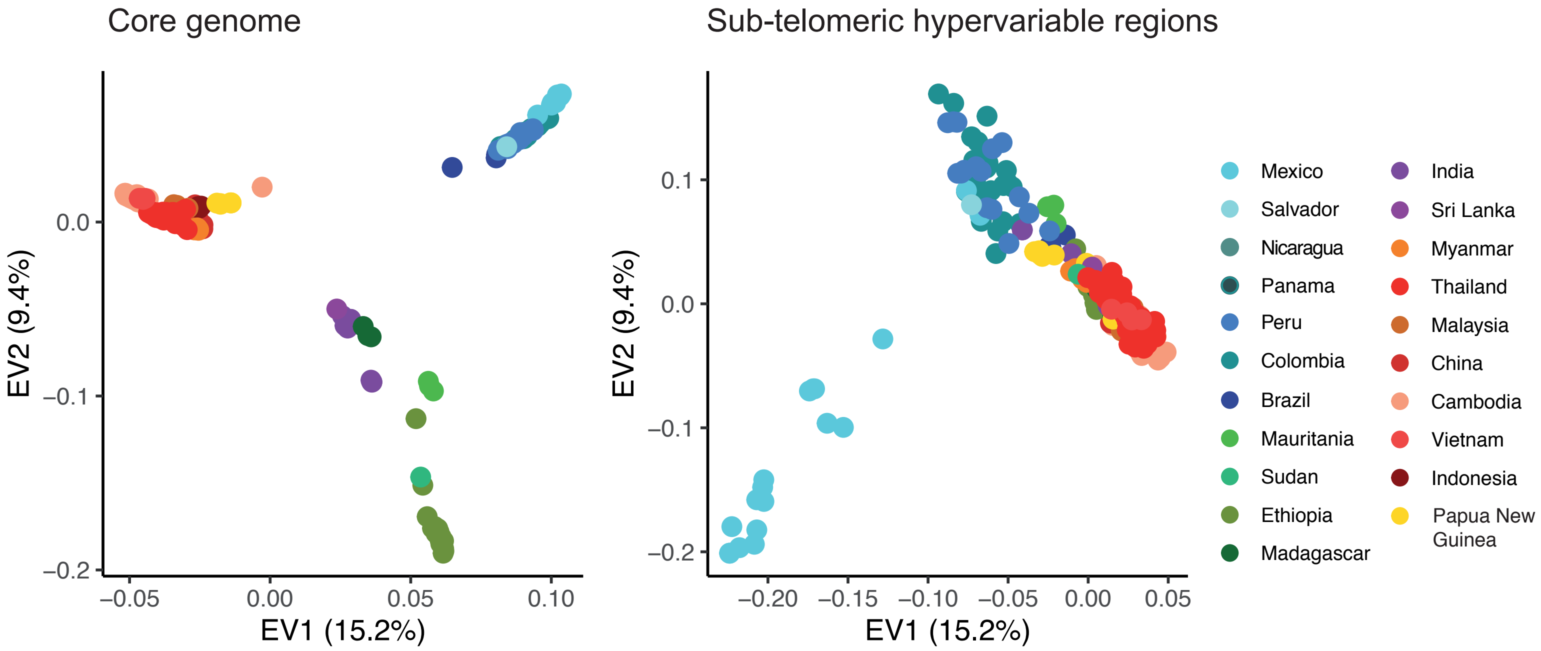


**Supplementary Figure 4:** Scatter plots showing the results of the two PCAs conducted using the SNPs located either in the core genome or in the hypervariable sub-telomeric regions.

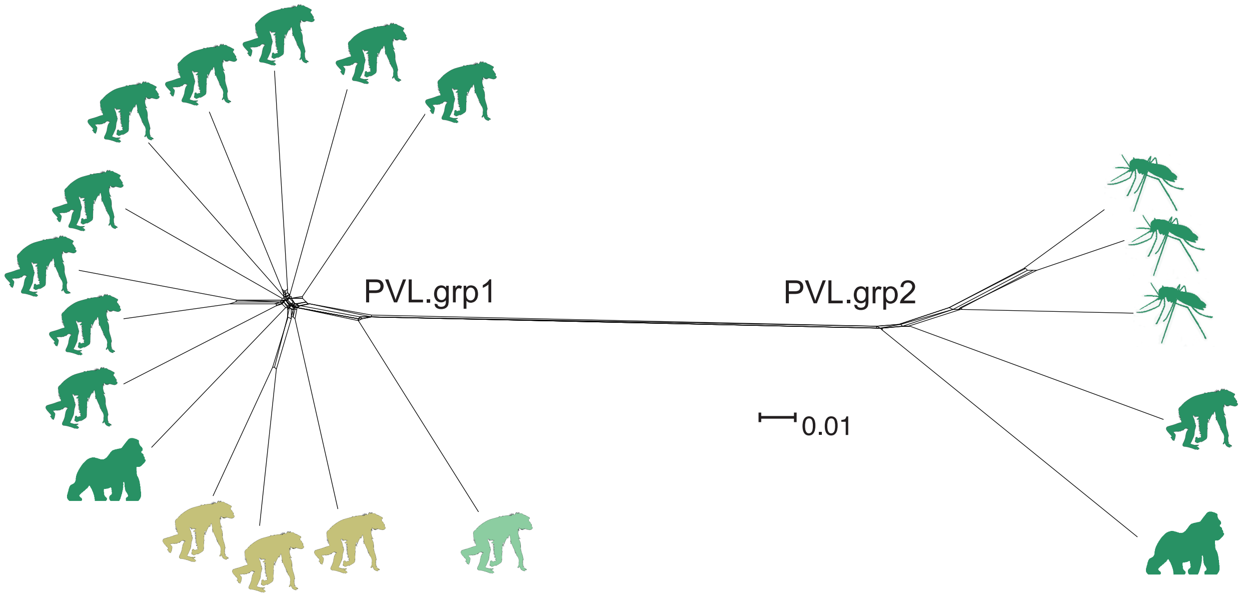


**Supplementary Figure 5:** Reticulate network based on SplitTree4 (ref), drawn from 19 *P. vivax-like* genomes. The reticulate networks split the P. vivax-like genomes in two distinct clades. On the tree, reticulation indicates likely occurrence of recombination. No reticulation was observed among the two clades suggesting the absence of recombination between the two clades.

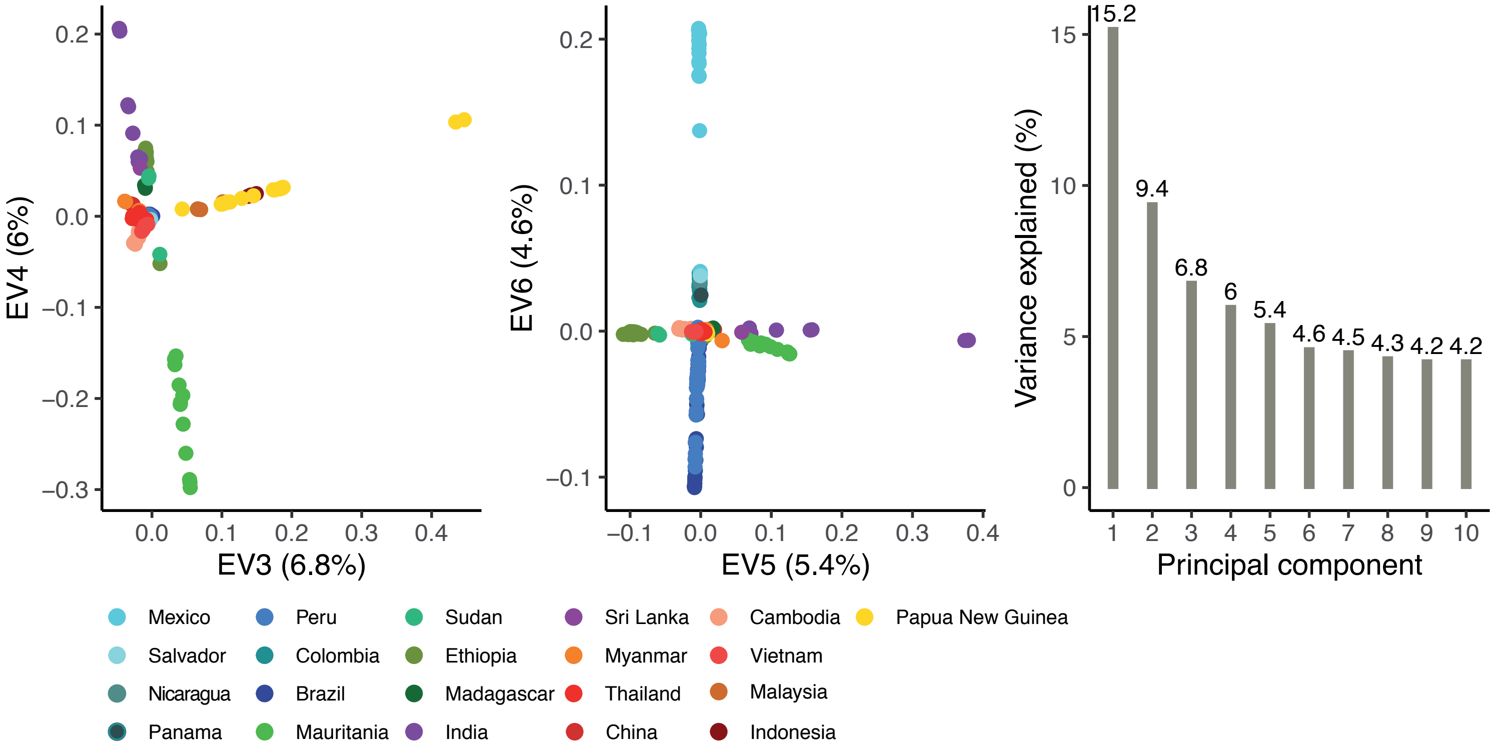


**Supplementary Figure 6:** PCA analysis of the 447 of *P. vivax* using SNPs present in the core genome. Scatter plot shows individual strain relationships along the principle components 3 to 6. Dot colors indicate populations. The bar chart shows the percentage of variance explained by each principal component axis.

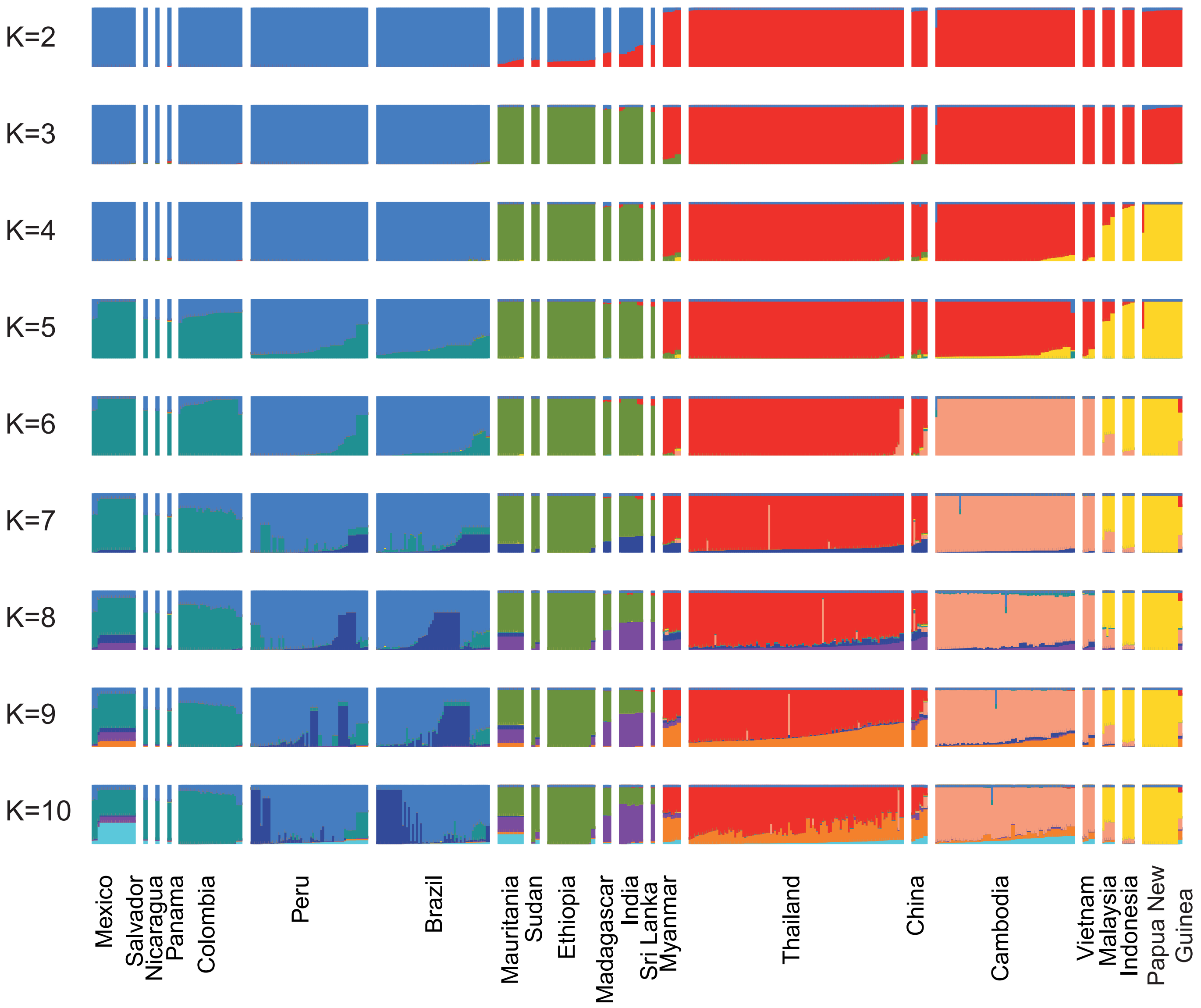


**Supplementary Figure 7:** Individual ancestry proportion to each ancestral population tested (from K=2 to K=10) estimated using the ADMIXTURE program for the 447 *P. vivax* genomes. Individual strains are sorted by country with increasing longitude.

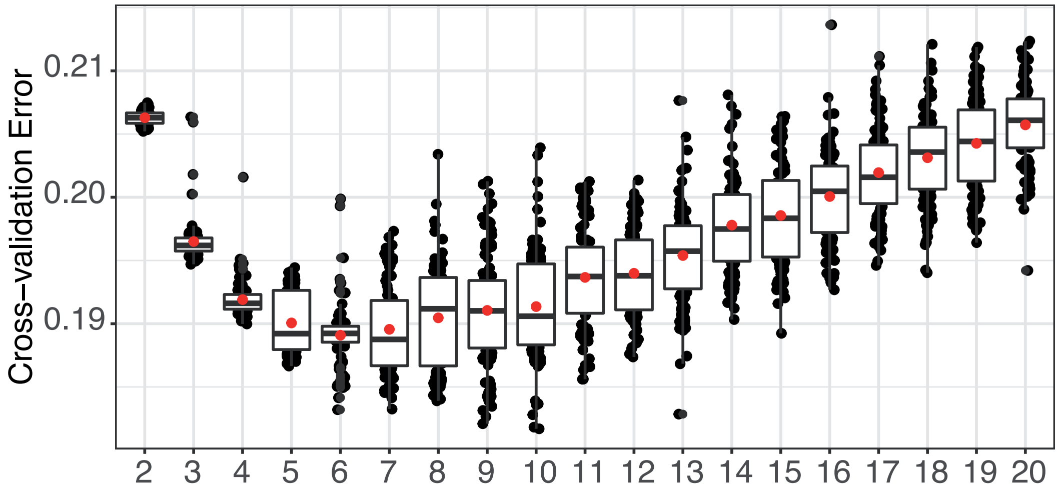


**Supplementary Figure 8:** Cross validation error rate estimated using the ADMIXTURE program for each ancestral populations tested (K between 2 and 20) .We chose K=5 to analyze the SNPs data, as the value that minimizes the error.

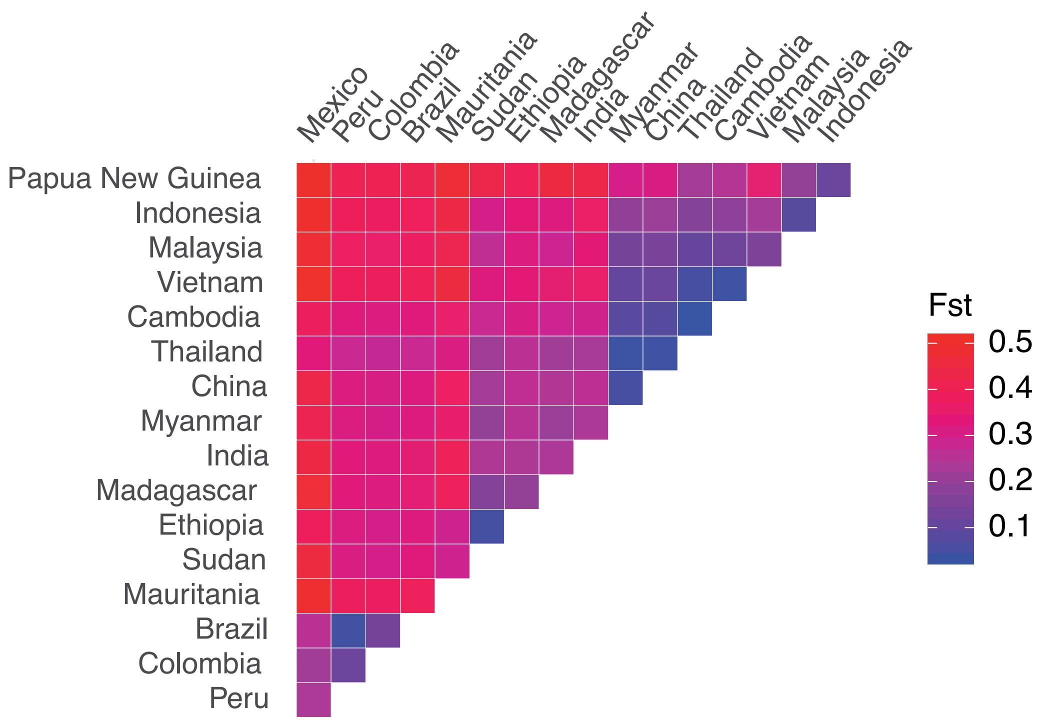


**Supplementary Figure 9:** Average allele frequency differentiation estimated using FST values between pairs of populations.

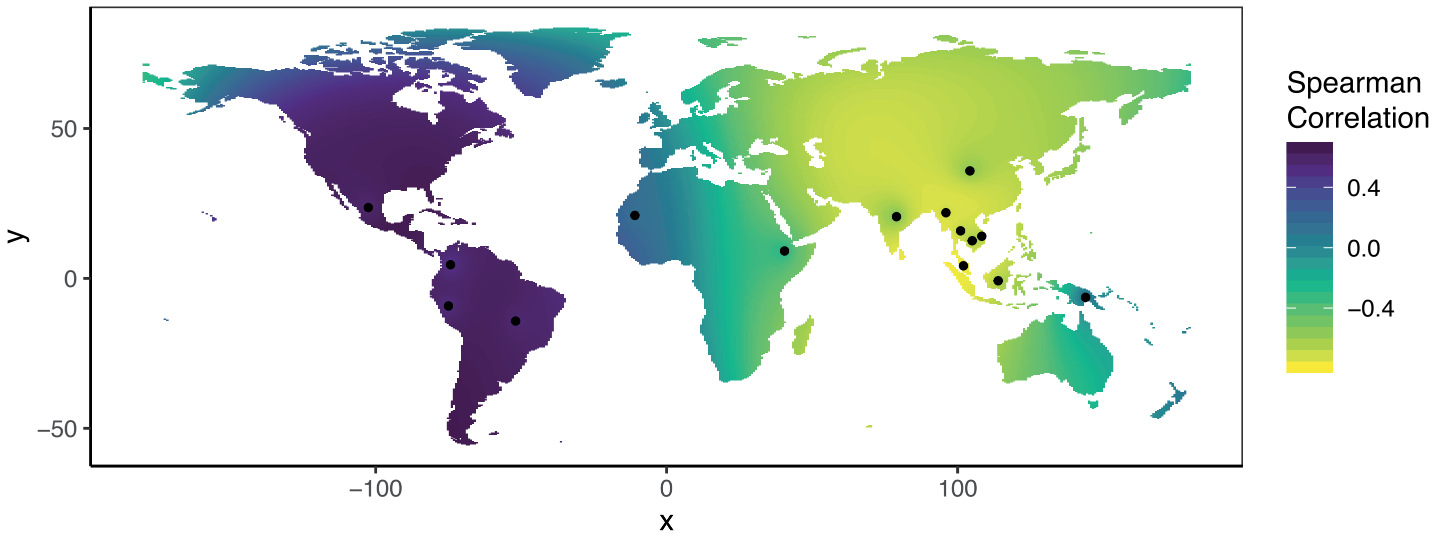


**Supplementary Figure 10:** Genetic diversity of *P. vivax* regressed on geographic distance across the world. The value at each pixel of the map corresponds to the Spearman correlation coefficient (r) between the expected genetic diversity in each population and the geographic distance between this population and the pixel. The black dots represent the sampling sites used in the regression (where n ≥ 5 individuals).

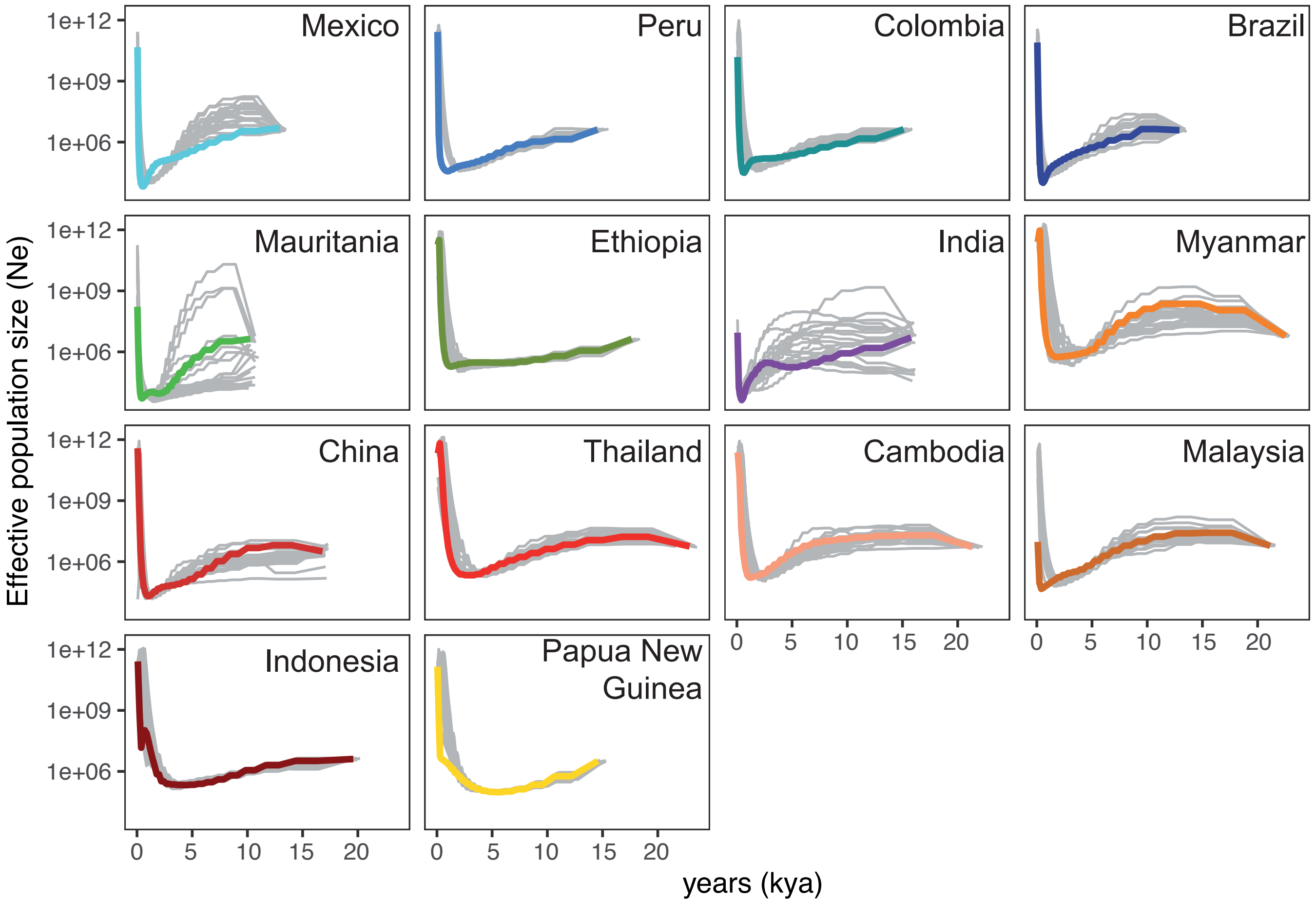


**Supplementary Figure 11:** Multiple sequentially Markovian coalescent (MSMC) estimates of the effective population size (Ne) in 14 *P. vivax* populations. The y-axis shows the log10 of Ne. Gray lines represent the MSMC results obtained from 50 bootstrap resampling of the segregating sites.
